# Antagonistic regulation with a unique setpoint, integral and double integral action

**DOI:** 10.1101/248682

**Authors:** Kristian Thorsen, Peter Ruoff, Tormod Drengstig

## Abstract

Several biochemical species are in organisms controlled in a pairwise manner i.e., two different species (e.g., hormone, enzyme, transporter protein) work to control the concentration of a third chemical species. Such pairs are often antagonistic, meaning that one of the controller species acts to increase whereas the other controller species acts to decrease the amount of the controlled species. How antagonistic systems interact to achieve regulation and to avoid competing against each other is not fully understood. An issue is how two antagonistic hormones can agree upon one common setpoint. We present here a new type of antagonistic regulatory system that has a single unique setpoint inherently defined by the system. The regulatory system controls the concentration of a chemical species with both integral and double integral action, achieving tight control. We show by the use of an analytical stability analysis, using the principle of vanishing perturbations, that the setpoint is asymptotically stable. Finally the prospect of treating the presented system as a part of a larger family of antagonistic regulatory systems with unique setpoints, integral and double integral action is discussed.

## 1. Introduction

Organisms have the property to adapt to a changing environment and keep certain components within the organism tightly regulated. In physiology and biochemistry this property is called homeostasis [1–4]. In control theoretic terms homeostasis is the ability of a system to self-regulate one or more of its states to a certain value under changing environmental conditions. The phenomenological laden biophysical concepts of homeostasis and adaptation [1, 3, 4] are closely related to mathematical concepts from control and systems theory [5–10].

Many homeostatic systems are regulated in a pairwise manner, where one chemical species acts to increase the concentration of a controlled species while another chemical species acts to decrease the very same concentration. Examples include the regulation of blood glucose by the antagonistic hormones insulin and glucagon [2], and blood calcium homeostasis by calcitonin and parathormone [2]. Another example is iron homeostasis in yeast where up-take (inow) is regulated through the protein complex Fet3p-Ftr1p which is induced by the iron sensing transcription factor Aft1p [11, 12]. The transcription factor Aft1p also activates Cth2p, which again specifically down-regulates proteins that participate in many Fe-dependent and consuming processes (outflow) [12, 13].

How the homeostatic value or the setpoint is achieved in antagonistic systems, and why hormones often appear in antagonistic pairs is not fully understood. An early theory is the so-called *rein control* introduced by Clynes [14]. Rein control is however, as pointed out in [15], not directly compatible with the idea of integral control and regulation toward a single setpoint with no steady state error. An integral controller is operating until the error between the controlled species and the setpoint is zero, and as stated by Saunders: ”If there are two independent integral controllers in a system, then unless the set points are precisely the same, which cannot be guaranteed in any real system, least of all in physiology, at least one of them (in practice generally both) will always be operating. Hence a proper steady state cannot be achieved even asymptotically” [15]. One way to achieve a single setpoint in a system with two integral controllers is to have them directly interact with each other. Saunders and Koeslag have shown how two controllers (in their case hormones) with a more or less symmetric mutual inhibition between them can provide a antagonistic system with a single setpoint, in what they call *integral rein control* [15, 16].

We have previously presented a set of two-component controller motifs with integral action[17]. These motifs can based on whether regulation is achieved by adjusting the inow or the outow of the controlled species be divided into what we call inow and outow controllers. We also showed how such inow and outow controller motifs can be combined to form antagonistic pairs exercising control on a common chemical species, and gave several physiological examples [17]. The combinations we focused on in [17] were all made out of two individual controllers and did hence not have a single *unique* setpoint. When the setpoint of the inflow controller motif is lower than the setpoint of the outflow controller motif, the steady state level of the controlled species will, dependent on the inflow or outflow perturbations, either end up at one of the setpoints or stay somewhere between them [17]. In that case both controllers cannot achieve zero steady-state error, between the regulated value and the setpoint, at the same time.

As a continuation of our previous work we present here a new theoretical model of an antagonistic biological regulatory system with a single unique setpoint. The system is formed by combining two single controller motifs in a nonindependent way. Specifically by having one of the motifs interact directly with the other, but without the interaction being mutual the other way around. This provides both integral and double integral action in the regulation of the controlled species. Double integral action is achieved by having one of the antagonistic controller species affecting the production or removal of the other.

## 2. Investigated System

Fig. 1A shows the reaction kinetic representation of the investigated system. The system consists of three chemical species *A*, *E*_1_ and *E*_2_, where *A* is held at the setpoint Aset by the two controller species *E*_1_ and *E*_2_. The controller species, *E*_1_ and *E*_2_, acts on compensatory flows; the synthesis/inflow of *A* is controlled by *E*_2_ and the degradation/outflow of *A* is controlled by *E*_1_. These flows compensate for other uncontrolled inflows and outflows of *A* which is represented by 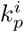 and 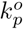 (disturbances).

**Figure 1:**
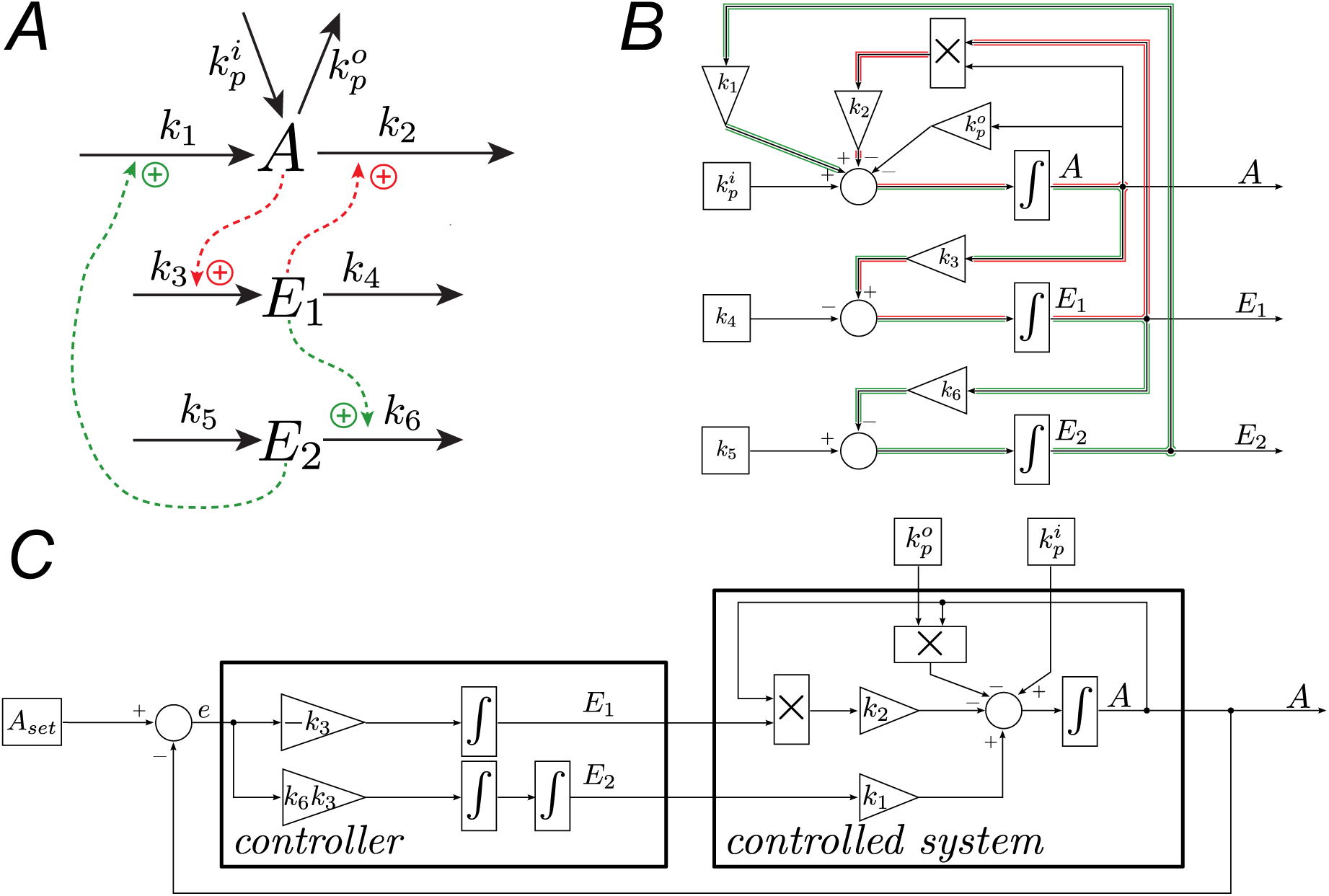
**A** Reaction kinetic representation of the system. *E*_1_ and *E*_2_ controls the outow and inow of *A*, respectively. Solid arrows represents mass ow and dotted arrows with plus signs represents biochemical activation. **B** Block schematic representation of the system. The integral and double integral feedback loops are highlighted in red and green, respectively. **C**. Block schematic representation showing the distinction between the controller and the controlled system. *E*_1_ and *E*_2_ acts as two inputs to the controlled system. The value of these inputs are proportional to the integral and the double integral of the error between the setpoint (*A*_*set*_ = *k*_4_/*k*_3_) and the current value of *A*, which is fed back from the controlled system to the controller.

Furthermore, *A* induces synthesis of *E*_1_, and *E*_1_ activates degradation of *E*_2_. A perturbation that increases the level of *A* (e.g., an increase in 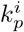 or a decrease in 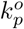) leads to increased synthesis of *E*_1_, which counters the perturbation by increasing the degradation of A. In addition the increased amount of *E*_1_ activates the degradation of *E*_2_, which again limits the inflow of *A*. As a whole these two mechanisms achieve tight control of *A* by adjusting both the synthesis and the degradation at the same time. A perturbation that decreases the level of *A* will have the opposite effect. The controller species *E*_1_ and *E*_2_ can be considered to function as biochemical outflow and inflow controllers [17].

The mathematical model of this system is found by applying the law of mass balance to derive differential equations for *A*, *E*_1_ and *E*_2_. In our system the inflow of *A* is proportional to the concentration of *E*_2_, expressed as *k*_1_*E*_2_. The outflow of *A* is dependent on the available amount of *A* and *E*_1_, expressed as *k*_2_*AE*_1_. The synthesis of *E*_1_ is taken to be proportional to the concentration of *A*, as *k*_3_*A*, and the synthesis of *E*_2_ occurs at a constant (basal) rate, *k*_5_. In order to achieve robust integral control of *A*, the degradation of the controller species *E*_1_ and *E*_2_ must follow zero-order kinetics in relation to their own concentrations [18]. As elucidated in [18], and more recently put in relation to synthetic biology by Ang and McMillen [19], such zero-order kinetics can be achieved in a biochemical environment if *E* is degraded by a saturated enzyme (e.g. Michaelis Menten kinetics with a low *K*_*M*_ value). The degradation of *E*_2_ is taken to be influenced by *E*_1_ in a linear manner.

To summarize, our 3rd order nonlinear model is described by the following differential equations:

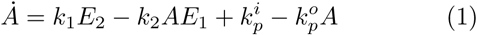

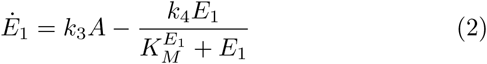

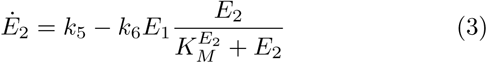

where *k*_*i*_ ∀ *i* = 1,…, 6, 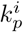, and 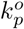 are positive rate constants.

Note that the goal here is to present this new motif and show how it works; the relative simple kinetic expressions for activation and removal can of course be replaced with more complex expressions that include features such as saturation, e.g., Hill functions. Refer to [20] and [21] for a treatment of such features affects the performance of the original two species controller motifs.

## 3. Integral and double integral action

This system has both integral and double integral action in the control of *A*. *E*_1_ accounts for the integral action, while *E*_2_ gives extra control by double integral action.

### 3.1. Integral action

To show that *E*_1_ works as an integral controller of *A* we begin by rewriting Eq. (2):

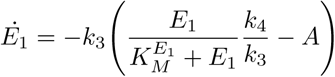

Assuming near zero-order removal of 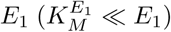, this expression reduces to

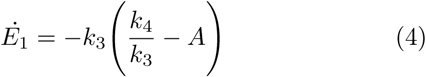

from where the steady state condition in *E*_1_ leads to the expression for the theoretical setpoint of *A* as *A*_*set*_ = *k*_4_/*k*_3_. By comparing Eq. (4) with the standard integral control law in its derivative form:

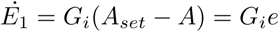

we see that *E*_1_ acts as an integrator that integrates the error *e* between the setpoint *A*_*set*_ and the actual value of *A*. The integral gain *G*_*i*_ = –*k*_3_ is negative, which is necessary as *E*_1_ is helping removing excess of *A*, i.e. an increase in *E*_1_ will decrease *A*. As will be seen below, it is helpful to write *E*_1_ on its integral form:

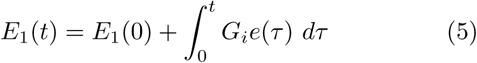

### 3.2. Double integral action

Whereas *E*_1_ acts as an integral controller, *E*_2_ acts as a controller with double integral action. To show this we start by rewriting Eq. (3), assuming 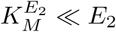

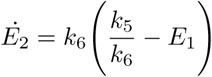

from where the steady state condition in *E*_2_ leads to the expression for the equilibrium point of *E*_1_ as 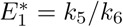. By inserting the integral expression for *E*_1_ from Eq. (5), we get

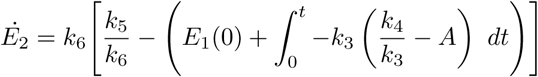

Suppose that *E*_1_(*t*) at the starting time *t*=0 is in its equilibrium, then this expression reduces to

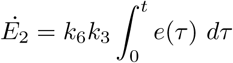

Then by writing *E*_2_ on its integral form we clearly see that *E*_2_ acts as a double integrator of the error between the setpoint and the actual value of *A*,

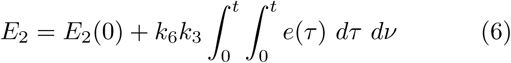

The system given by the differential equations in Eqs. (1), (2), and (3) can be represented in a block schematic form, shown in Fig. 1B. The first feedback loop with single integral action can be identified by following the line from *A* via *k*_3_, the *E*_1_ integrator, the multiplication block, and *k*_2_, colored red in Fig. 1B. By dividing *k*_4_ with *k*_3_ the *k*_3_ gain block can be moved to after the summation, showing more clearly how *k*_3_ acts as the integral gain parameter of the *E*_1_ controller.

Instead of following *E*_1_ directly back to *A*, the second feedback loop with double integral action can be identified by following *E*_1_ via *k*_6_ and the *E*_2_ integrator back to *A* through *k*_1_, colored green in Fig. 1B.

The block schematic form shown in Fig. 1C simplifies the analysis by arranging the blocks in such a way that the two controllers, given by Eqs. (5) and (6), are clearly shown in a feedback loop, controlling the controlled system. The two controllers compensate for disturbances, i.e. 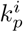 and 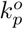, affecting *A*.

## 4. Stability Analysis

We will here show that the presented regulatory system is stable. We will first show that the system is stable in some local region about the origin by using linearization, and then broaden this region by using the principle of vanishing perturbations. In the analysis we assume that *E*_1_ and *E*_2_ are degraded by perfect zero-order mechanisms, i.e. 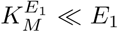 and 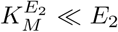, so that the simplified system equations can be written as:

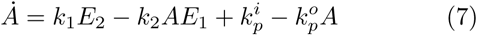

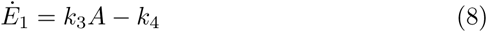

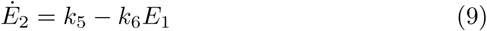

Since negative concentrations are unphysical our states are restricted to non-negative values. However, the simplified system is not always non-negative, due to the use of zero-order terms instead of Michaelis Menten kinetics. Hence, it is only a good enough representation of the original system as long as the trajectories stays some distance away from zero-values. This is equivalent to the conditions 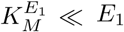 and 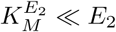 being true for all encountered values of *E*_1_ and *E*_2_.

From Eqs. (7), (8) and (9), we find the equilibrium point to be

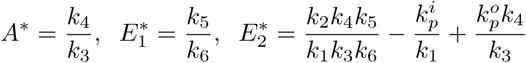

To simplify the following stability analysis, we want to move the equilibrium point to the origin by doing a change of variables. It is fairly simple to show that the transformed system, with an equilibrium point at the origin, can be written as:

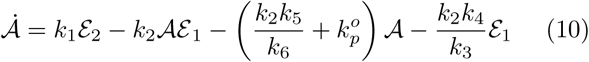

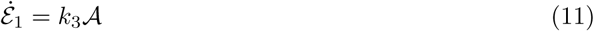

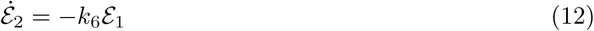

Where 𝒜, 𝓔_1_ and 𝓔_2_ are the transformed states. This system will also be referred to by the notation *ẋ* = *f*(*x*), where *x* = [𝒜, 𝓔_1_, 𝓔_2_]^*T*^.

### 4.1 Local stability

We use Lyapunov’s indirect method, i.e. linearization, to check for local stability. The Jacobian matrix evaluated at the origin is,

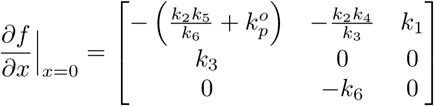

where eigenvalues can be found from the characteristic equation:

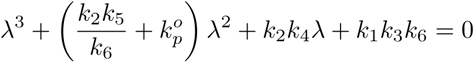

The Routh-Hurwitz criterion [22] can be used to provide the following necessary and sufficient condition for the equilibrium point to be locally asymptotically stable

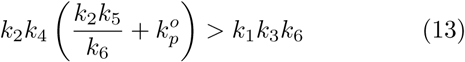

### 4.2 Stability by the principle of vanishing perturbations

To use the principle of vanishing perturbations as given in lemma 1 in the appendix, we treat the linearized version of the perturbed system evaluated at the origin, as the nominal system. The perturbed system is written as a nominal linear system Φ*x* perturbed by a nonlinearity *ϕ*(*x*) that vanishes at the origin as

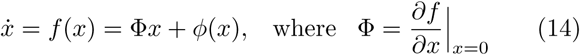

and *ϕ*(*x*) is the remaining part necessary for the equality in Eqs. (14) to hold.

The nominal system *ẋ* = Φ*x* is asymptotically stable when the condition in Eqs. (13) is fulfilled. For a asymptotically stable system there exists a solution to the Lyapunov equation [23]

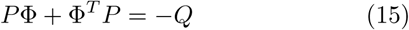

where *P* and *Q* are positive definite symmetric matrices. From [23] we have that the quadratic Lyapunov function *V*(*x*) = *x*^*T*^*Px* satisfies

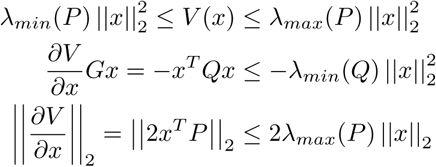

where || · ||_2_ is the Euclidean norm, and *λ_min_* and *λ_max_* are the minimum and maximum eigenvalues. This fulfills the first three conditions in lemma 1, (A.2), (A.3) and (A.4), with the constants *c*_1_ to *c*_4_ given by:

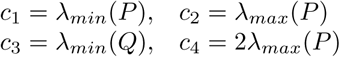

The perturbation consists of the nonlinear cross term from Eq. (10)

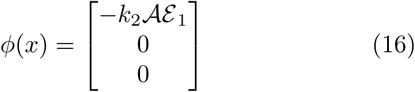

Using the quadratic function *V*(*x*) = *x*^*T*^*Px* as a candidate Lyapunov function for the perturbed system, the derivative along the trajectories is

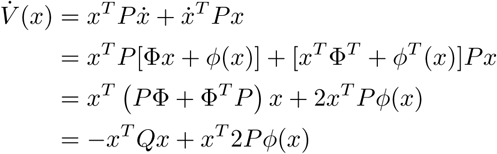

Looking at the perturbation term in Eq. (16), we only have to limit *ϕ*_1_(*x*), i.e. the 1st element in *ϕ*(*x*). We would prefer to limit 𝒜 instead of 𝓔_1_, since this will give a result that says something about how far perturbations can drive 𝒜 from its setpoint for the regulatory system to still be able to regulate 𝒜 back to the setpoint. Hence, we limit 𝒜 with

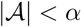

and we can now limit *ϕ*_1_(*x*) and *ϕ*(*x*) by the bound

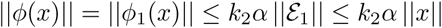

From the principle of vanishing perturbations (lemma 1 in the appendix) the origin of the whole perturbed system is exponentially stable if:

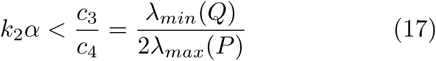

Using the identity matrix as *Q*, i.e. *Q* = *I*, will maximize the ratio λ_*min*_(*Q*)/2λ_*max*_(*P*) [23].

To sum this up our system is stable when:

1. The nominal system is asymptotically stable, which is true when the Routh-Hurwitz based criterion in Eq. (13) holds. Since the nominal and the linearized perturbed system are the same this means that the whole perturbed system has to be locally asymptotically stable.
2. Then we can always find an limit on |𝒜| < *α* so that the condition in Eq. (17) holds, and the whole system will be asymptotically stable in the region defined by |𝒜| < *α*.

## 5. Numerical Example

We illustrate the properties of the system by considering the case when the constants *k*_1_ to *k*_6_ are [1, 1, 1, 2, 2, 0.5] 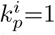=1, and 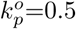=0.5, all in arbitrary units (a.u.). The equilibrium point of the original system Eqs. (7), (8) and (9) is

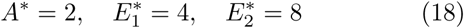

### 5.1. Regulatory action

The ability to keep *A* at its setpoint *A*_*set*_ = *k*_4_/*k*_3_ is demonstrated by the simulation shown in Fig. 2A. We perturbed *k*_4_ to make the setpoint shift in a stepwise manner and then perturbed *k*_4_ linearly so that the setpoint changed as a ramp function. A key property of regulatory systems with double integral action is their ability to follow a ramping setpoint [24]. The simulation also shows that *A* is the only variable that is regulated exactly to its setpoint during the ramping phase. The equilibrium of *E*_1_ is independent of *k*_4_, and hence, does not change in the simulation, see the dashed red line in Fig. 2B. Still, the system shows adaptation in the level *E*_1_ during step changes in the setpoint of *A*, but not during ramping as shown in Fig. 2B. The equilibrium of *E*_2_ is changed by the perturbations in *k*_4_. However, *E*_2_ is not homeostatic regulated during the ramping phase, see Fig. 2C.

**Figure 2:**
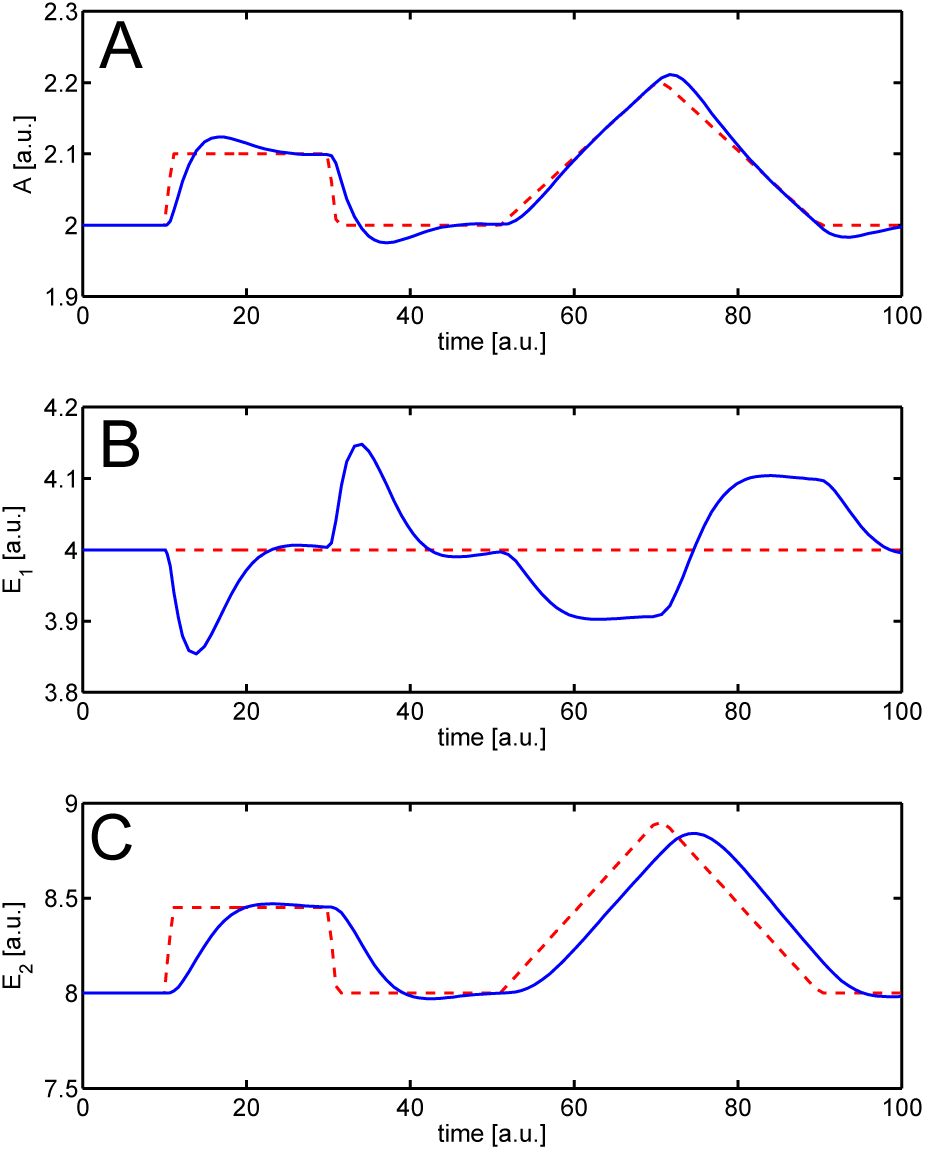
Response in *A*, *E*_1_ and *E*_2_ (Eqs. (7)–(9)) during setpoint changes induced by perturbing *k*_4_ to 2.1 during *t* = {10, 30}, increasing it linearly from 2.0 to 2.2 during *t* = {50, 70} and then decreasing it linearly from 2.2 to 2.0 during *t* = {70, 90}. The rest of the parameters are set to *k_i_* = [1, 1, 1, 2, 2, 0.5], 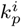=1, and 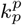=0.5. Panel **A**, **B** and **C** show the level of *A*, *E*_1_ and *E*_2_ (solid blue) together with their respective equilibrium points (dashed red).

To illustrate the disturbance rejection capabilities of the regulatory system we changed the uncontrolled inflow and outflow by perturbing 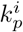 and 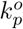. The results show perfect adaptation/homeostatic regulation of *A* for both stepwise and ramping disturbances, see Fig. 3.

**Figure 3:**
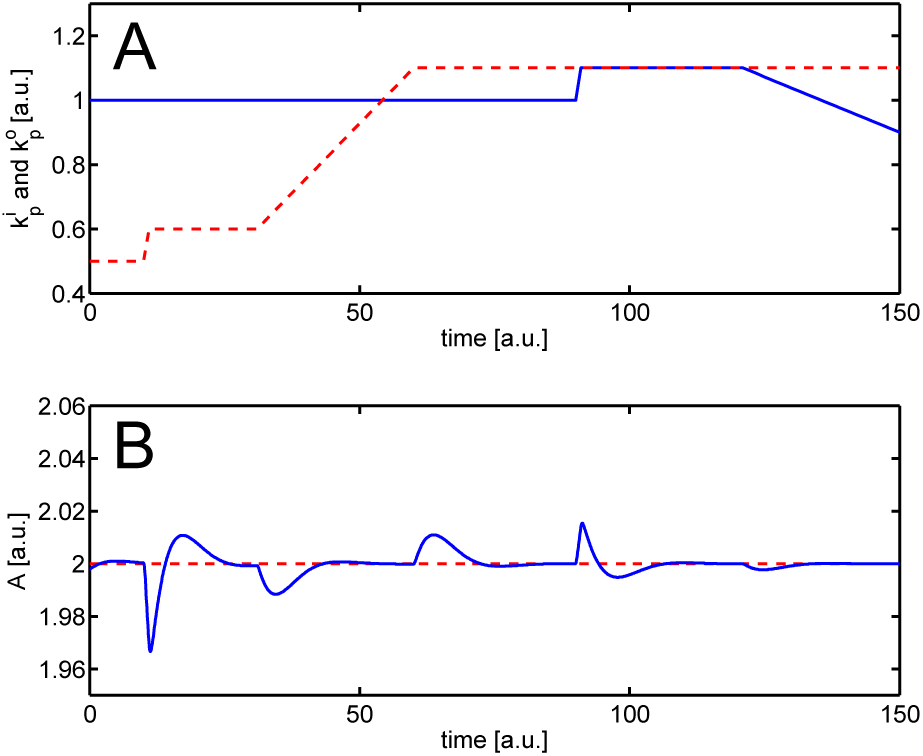
Response in *A* (Eqs. (7)–(9)) during perturbations in the uncontrolled inflow and outflow. The parameters are set to *k*_*i*_ = [1, 1, 1, 2, 2, 0.5], 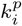 (solid blue) and 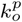 (dashed red) are changed as shown in panel **A**. Panel **B** shows the level of *A* (solid blue) together with its setpoint (dashed red). Due to the double integral action *A* adapts not only to step-type disturbances (*t* = {10, 30} and *t* = {90,120}) but also to ramp-type disturbances (*t* = {30, 60} and *t* = {120, 150}).

### 5.2 Stability

By using the rate constants above, condition Eq. (13) holds, and the system is locally stable. The Jacobian matrix evaluated at the origin is:

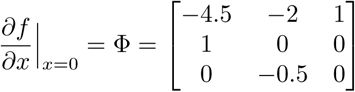

Since the system fulfills the criterion in Eq. (13), we know that the system is asymptotically stable in some neighborhood around the origin. The size of this neighborhood can, however, not be quantified directly by Lyapunov’s indirect method. Using *Q* = *I*, we solve the Lyapunov equation Eq. (15), and find *P* to be

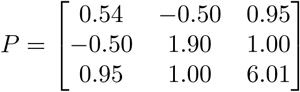

which has the eigenvalues 0.12, 1.90, and 6.36. Using the largest eigenvalue of *P*, we find the bound *α* from Eq. (17) as:

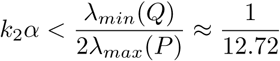

Using *k*_2_ = 1 gives *α* < 0.078. The region |𝒜| < 0.078 translates to *A* ∈ (*A*_*set*_–0.078, *A*_*set*_+0.078). So the system is guaranteed to be asymptotically stable as long as *A* is between 1.922 and 2.078. This bound may seem strict, but the analysis does not say that the system is unstable outside this region, only that the analysis as done cannot prove stability outside this region.

The perturbation term is in the description of the principle of vanishing perturbations (lemma 1 in the appendix) given as a term that can affect all of the states directly. In our system however, the perturbation term does only directly affect the derivative of *A*, see Eq. (16). This fact can be used to improve the analysis. First we examine the derivative of the Lyapunov function along the system trajectories, inserting the perturbation term *ϕ*(*x*) and the *P* matrix:

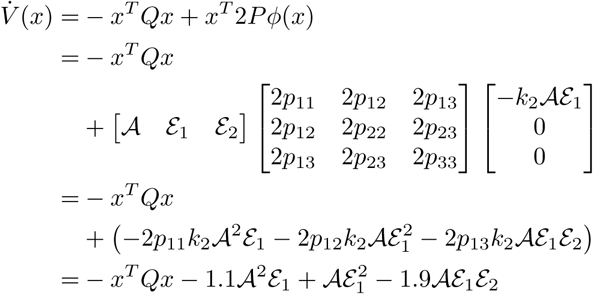

where we have inserted the values for *k*_2_ and the elements in the *P* matrix. By limiting *A* with |𝒜| < *α*, we can limit *V̇*(*x*) by:

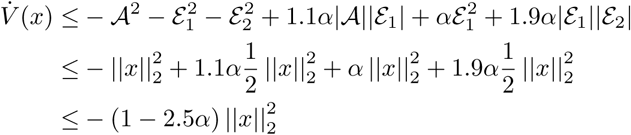

and *V̇* (*x*) is negative definite when *α* < 0.4. The system is then asymptotically stable by Lyapunov’s direct method as *V*(*x*) is guaranteed to be positive definite by the positive definiteness of the *P* matrix. This result give us a bound on 𝒜 which is less strict than what was given by the preceding analysis. The region |𝒜| < 0.4 translates to *A* ∈ (*A*_*set*_ – 0.4, *A*_*set*_ + 0.4), so the system is guaranteed to be asymptotically stable as long as *A* is between 1.6 and 2.4. Fig. 4 shows the phase plane and the trajectories of the system; it shows that numerically the trajectories tends to the equilibrium point inside as well as outside of the stability region found from this latter analysis.

**Figure 4:**
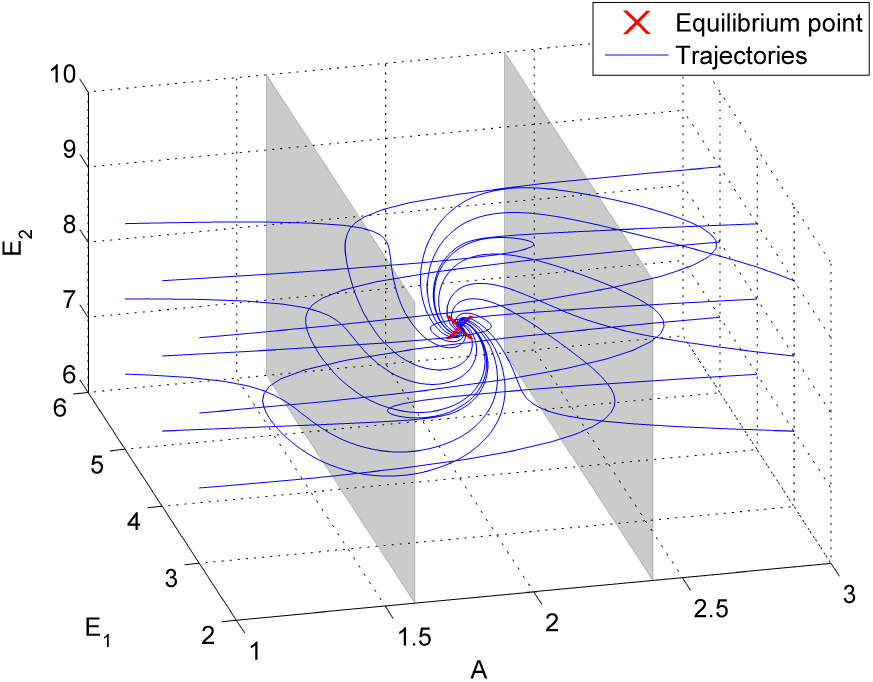
Phase plot for the system in Eqs. (7), (8) and (9) where the constants *k*_1_ to *k*_6_ are [1, 1, 1, 2, 2, 0.5]. The equilibrium point is marked as a red ’x’, and corresponds to Eq. (18). Trajectories starting from different initial conditions converges to the equilibrium point. The system shows stability also outside the region *A* ∈ (*A*_*set*_–0.4, *A*_*set*_ + 0.4).

## 6. Discussion and Conclusion

### 6.1. Regulatory properties of double integral action

From control theory it is well known that a control system made to perfectly follow, i.e. with no steady-state error, a parabolic reference of order *m* must, at least, have *m* +1 integrators in the forward loop [24]. This explains why a ramping reference, which is a parabolic of order 1, can be perfectly followed by the presented system, as shown in Fig. 2.

Although the ability to follow a reference is of primary interest in man made engineering systems, it may be of less importance in biophysical systems (discussed later). Adaptation, the ability to reject disturbance signals while keeping the output at a fixed setpoint is, on the other hand, often of prime interest. An alternative way to state the principle of integrators needed to follow a parabolic input is by which class of disturbances a regulatory system is able to regulate against, i.e., perfectly adapting to. This is described mathematically by the internal model principle (IMP) [25–27]. It states that in order for a regulatory system to asymptotically adapt its output *y*(*t*), under perturbations by a specified class of external disturbance signals, the system must necessarily contain a subsystem which itself is capable of creating the same class of signals, and this subsystem must do so while having only the system output *y*(*t*) as its input. In other words, a model of the disturbance signals must be present internally in the system. In our case this subsystem is the controller; a system with two integrators is able to produce ramping functions from a constant input *y*(*t*) (the system output *y*(*t*) is in the case of perfect adaption constant in its steady state).

We have in essence, by arranging our original system (which we may denote ∑ as in [27]) as a controlled system (which we may denote ∑_0_ as in [27]) and a controller (which we may denote ∑_*IM*_ as in [27]), employed the internal model principle. Refer to Fig. 1B and Fig. 1C.

It must be mentioned that *E*_2_ is also ramping during a ramping disturbance, which is exactly why *A* can stay constant. Since *E*_2_ in a real biological system can neither decrease below zero or increase to infinity, there are limitations, and the three-component controller motif will in such cases break down.

### 6.2. A class of antagonistic systems

The three-component structure used in this work can be extended to a whole class, or family, of three-component antagonistic regulatory systems. Fig. 5A shows the basic reaction kinetic structure for this class of systems. The controlled species *A* is antagonistic regulated by two interacting regulatory species *E*_1_ and *E*_2_; by antagonistic we hold that one of the two regulatory species acts on synthesis/inflow of *A*, while the other acts on degradation/outflow of *A*.

**Figure 5:**
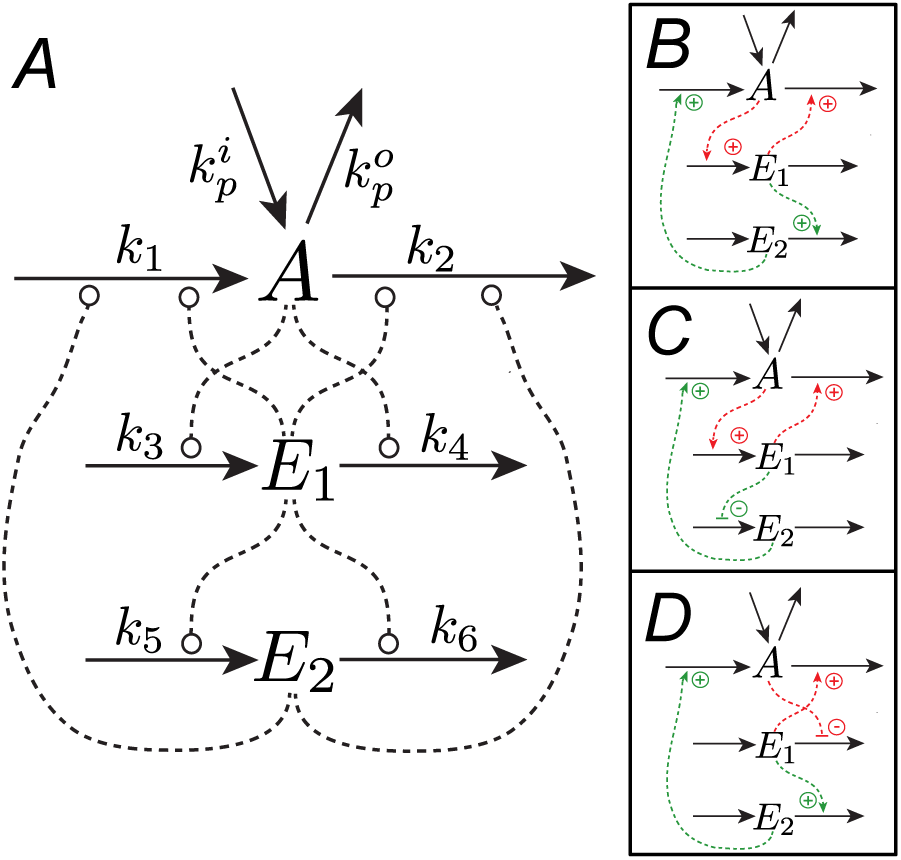
Panel **A** shows the basic structure of a three component antagonistic controller motif. Dotted lines show possible interactions, which can either be activating or inhibiting (round symbols). Not all interactions are allowed at the same time, see main text for explanation of the possible connections. Panels **B** to **D** shows three of the 32 possible configurations with negative *inner loop* and negative *outer loop.*

Taking that *A* can either act on the inflow or outflow of *E*_1_ by either activation or inhibition, and that *E*_1_ can affect *A* in the same manner we can construct a total of 16 different *inner loops* between *A* and *E*_1_. These *inner loops* are structurally the same as the 16 two-component controller motifs we presented in [17, Suppl.]. Half of these have negative feedback and the other half have positive feedback. Letting *E*_1_ act on either the inflow or the outflow of *E*_2_ by either activation or inhibition, we extend the number of possible configurations to 64. This number is further extended to 128 given that *E*_2_ must affect *A* by either activation or inhibition of the flow *not* affected by *E*_1_, completing the *outer loop*, and the requirement for antagonistic regulation. If we, however, let *E*_1_ and *E*_2_ act on the same compensatory flow, i.e., both acting on the inflow of *A* or both acting on the outflow of *A*, the total number of possible configurations is increased to 256. The last 128 configurations will not be antagonistic as we defined it above.

The integral action of the *inner loop* is present in all of the 8 configurations with negative feedback [17]. The double integral action achieved in our system requires an interaction between the two regulatory species, in a way such that the concentration of the second controller species *E*_2_ is dependent on the concentration of the first controller *E*_1_. Whether this dependence is positively coupled or negatively coupled, the latter being the case in our system, is not of primary importance, as the total action depends on how the second controller species acts on the controlled species *A*. The central point is that the *A* induced change in *E*_1_ counteracts the change in *A* not only directly, but also indirectly through *E*_2_. Hence, there will be 4 possible configurations of the *outer loop* for each of the 8 stable configurations of the *inner loop*, making a total for 32 possible configurations for three-component antagonistic systems with integral and double integral feedback.

Fig. 5 panels B to D show three of the 32 configurations. The configuration in panel B is the system described earlier in this work; the configuration in panel C is a similar system where the *outer loop* is changed by having *E*_1_ inhibit the inflow of *E*_2_ instead of activating the outflow. Panel D shows a configuration similar to panel B, but with an altered *inner loop*; A inhibits the outflow of *E*_1_ instead of activating the inflow. We are currently working on elucidating differences and individual properties within the class of three-component antagonistic regulatory systems, and plan to present the whole class in more detail at a later time.

### 6.3. Biological significance

One example where interactions between regulatory molecules (e.g. hormones) can be identified is the glucose, insulin, and glucagon system. Blood glucose is regulated by the two antagonistic hormones insulin and glucagon. Insulin is produced by the *β*-cells in the pancreas in response to high concentration of blood glucose. Insulin reduces the blood glucose by increasing the uptake of glucose by skeletal muscles and the liver. Glucagon is produced by the *α*-cells in the pancreas in response to low concentration of blood glucose. It works opposite to insulin by increasing the release of glucose from the liver, promoting gluconeogenesis and glycogenolysis [2]. The secretion of glucagon is dependent on blood glucose levels; more recent studies have, however, also revealed that insulin does affect the release of glucagon [28–30]. The *α*-cells are located downstream from the *β*-cells, making them highly exposed to the secreted insulin [31]. The presence of insulin has been shown to be essential for the suppression of glucagon secretion [30] and studies done in insulin receptor knockout mice have provided direct in vivo evidence on how insulin modulates the secretion of glucagon [28, 29]. This show that the required interaction between regulatory mechanisms to enable integral and double integral action as shown by our system is present, and may play a role in the regulation of blood glucose.

Instead of being mediated by some upstream or external factor the setpoint in the presented class of antagonistic systems is inherited in the structure, given by kinetic parameters. In this way, these three-component motifs are similar to the two-components motifs we have earlier presented [17]. The kinetic parameters are dependent on the spatial structure in which the biochemical processes occurs and the enzymes, cofactors, and transport proteins involved in the process. Since animals within a species usually all express the same proteins, this inherent setpoint can explain the similarity in defended levels among individuals [32].

Rheostasis, the condition in which the setpoint of a functioning regulatory system is changed over time, such as during certain phases of the life cycle, or as the response to an stimuli, such as the increase of body temperature (fever) during an infection [33], is one phenomenon that can be explained by such inherent definition of the setpoint. The setpoint may be changed by something as simple as increasing the level of an enzyme which catalyzes one of the reaction steps in the system.

Also, in relation to rheostasis, the double integral action can ensure that the regulated species is kept exactly at the setpoint when this is ramped up and down over time. Such close regulation can, however, be argued to be of little importance as no biological setpoint will be ramping forever. Also, some setpoint changes happens over such a long time that the regulatory system can be considered to be at steady state, this includes life cycle variations (programmed rheostasis), where the setpoint typically change over several days [33]. The discovery of such double integral action is still appealing.

Although real biological systems are necessarily more complex, with many more interacting substances and variables than our 3-component system, we have conceptually shown how antagonistic control can have both integral and double integral action and regulate a controlled species toward one inherently defined *unique* setpoint.

## A. Appendix

The appendix serves as a short reference for readers that are unfamiliar with principle of vanishing perturbations. The lemma also can be found in many textbooks on differential equations and stability analysis of dynamical systems, e.g. [23] or [34]. Notice that the perturbation term in the analysis should not be confused with uncontrolled perturbing inflow 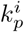 and outflow 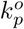 of *A*.

### Lemma 1. Principle of vanishing perturbations

*The following lemma considers stability for the resulting system when an exponentially stable nominal system with an equilibrium point at the origin is perturbed by a perturbation that vanishes (is zero) at the origin. Consider the system*

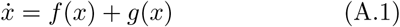

*where f*: 𝔻 → ℝ^n^ *and g*: 𝔻 → ℝ*^n^ are locally Lipschitz*^1^ *in x on* 𝔻, *and* 𝔻 ∈ ℝ^*n*^ *is a domain that contains the origin. The nominal system ẋ = f (x) is exponentially stable, and the perturbation satisfies g(x)|_x=0_ = 0. Let x = 0 be an exponentially stable equilibrium point of the nominal system. Let V(x) be a Lyapunov function of the nominal system satisfying the following conditions*

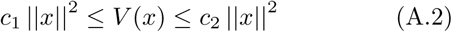

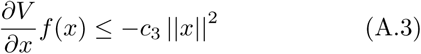

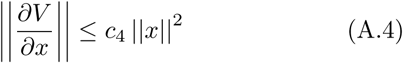

∀*_x_* ∈ 𝔻. *Suppose that the perturbation term satisfies the linear growth bound*

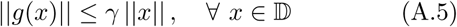

*and that*

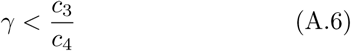

*Then, the origin is an exponentially stable equilibrium point for the whole perturbed system. For Proof see [23, chap. 9.1]*

## Individual contributions

Kristian Thorsen: initial idea, modeling, analytic and numerical studies, biological significance and discussion, drafted the manuscript, edited and revised the manuscript.

Peter Ruoff: proposed extending the presented system to a whole class, biological significance and discussion, edited and revised the manuscript.

Tormod Drengstig: modeling, proposed the presentation of regulatory action as done in Figs. 2 and 3, edited and revised the manuscript.

All authors have approved the manuscript.

## Footnotes

1 Lipschitz is a property weaker than continuous differentiability but stronger than continuity, it ensures uniqueness and existence of solutions to differential equations [23]

